# Structural studies of a serum amyloid A octamer that is primed to scaffold lipid nanodiscs

**DOI:** 10.1101/2023.12.08.570821

**Authors:** Asal Nady, Sean E. Reichheld, Simon Sharpe

## Abstract

Serum amyloid A (SAA) is a highly conserved acute-phase protein that acts on multiple pro-inflammatory pathways during the inflammatory response and is used as a biomarker of inflammation. It has also been linked to beneficial roles in tissue repair through improved clearance of lipids and cholesterol. In patients with chronic inflammatory diseases, elevated SAA may contribute to increased severity of the underlying condition. The majority of circulating SAA is bound to high-density lipoprotein (HDL), stabilizing SAA and altering its functional properties, likely through altered accessibility of protein-protein interaction sites on SAA. While high-resolution structures for lipid-free forms of SAA have been reported, their relationship with the lipid or HDL-bound forms of the protein, has not been established. We used multiple biophysical techniques, including SAXS, TEM, SEC-MALS, native gel electrophoresis, glutaraldehyde crosslinking, and trypsin digestion to characterize the lipid-free and lipid-bound forms of SAA. SAXS and TEM data show the presence of soluble octamers of SAA with structural similarity to the ring-like structures reported for lipid-free ApoA-I. These SAA octamers represent a previously uncharacterized structure for lipid-free SAA and are capable of scaffolding lipid nanodiscs with similar morphology to those formed by ApoA-I. SAA-lipid nanodiscs contain four SAA molecules and have similar exterior dimensions as the lipid-free SAA octamer, suggesting that relatively few conformational rearrangements are required for lipid binding. This study suggests a new model for SAA-lipid interactions and provides new insight into the ability of SAA to stabilize protein-lipid nanodiscs or even replace ApoA-I on HDL particles during inflammation.

## Introduction

Serum amyloid A (SAA) proteins are small apolipoproteins that are highly upregulated during the acute phase inflammatory response (1–3). Due to the increased levels of circulating SAA produced in response to inflammation, this protein has been used as a biomarker for a wide range of disease states (4–7). SAA proteins are highly conserved from echinoderm to humans, and an increasing number of studies suggest roles for SAA in both the acute inflammatory response, as part of the innate immune system and in altered lipid or cholesterol transport (8–11). SAA is also highly upregulated in patients with chronic inflammatory conditions, including autoimmune diseases, diabetes, cardiovascular disease, inflammatory bowel disease and COVID-19 associated hyper-inflammatory syndrome. In these instances, both hepatic and non-hepatic expression of SAA are considered to stimulate pro-inflammatory pathways and contributes to disease progression (12–16).

During the acute phase response, hepatic expression of SAA is upregulated up to 1000-fold, resulting in serum concentrations of ~1 mg/ml. The majority of this circulating SAA is bound to high-density lipoprotein (HDL), with low-density lipoprotein (LDL) and very low-density lipoprotein (VLDL) binding also observed to lesser degrees (17–20). SAA has been shown to displace some of the primary protein component of HDL, apolipoprotein A-I (ApoA-I), during acute inflammation, resulting in altered HDL functionality (21, 22). The presence of SAA has been shown to reduce the normal anti-inflammatory activity of HDL (23, 24). This leads to the activation of proinflammatory and proatherogenic pathways through increased interactions with toll-like receptors 2, and 4 (TLR-2 and TLR-4) to promote the release of proinflammatory cytokines, FPRL-1 to promote chemotaxis towards the site of infection/injury, and CD36 to promote lipid uptake (16, 22, 25, 26). SAA also alters the reverse cholesterol and lipid transport activities of HDL, further contributing to the development and progression of atherosclerosis and other cardiovascular diseases (27–29).

In addition to the pro-inflammatory activities of SAA, this protein has also been shown to play a role in the clearance of lipids and cholesterol from sites of tissue damage, facilitating repair and recovery (15, 30, 31). One proposed mechanism involves SAA binding to scavenger receptors and promoting cholesterol efflux (32). However, the reported ability of lipid-free SAA to recruit phospholipids into nanodiscs, even in the absence of ApoA-I, strongly suggests a more direct protective role for this protein in lipid clearance and highlights the importance of understanding the molecular level interactions between SAA and lipids (33–35).

There are four primary isoforms of SAA, each of which may also exhibit some allelic variation. SAA1 and SAA2 share 92% sequence identity and are both highly upregulated during the acute phase inflammatory response, SAA3 is a pseudogene in humans and the constitutively expressed SAA4 shares less sequence identity with SAA1 and SAA2 (25, 36). SAA1 and SAA2 are small 12 kDa apolipoproteins (106 residues) composed of 4 alpha helices followed by a largely disordered C-terminus (37, 38). Note that due to the hydrophobicity of helices 1 and 3, the SAA monomer is poorly soluble, such that the two crystal structures of SAA solved to date portray a similar monomer structure (cone shaped 4-helix bundle) in the context of two different oligomeric states (hexameric SAA1 vs. tetrameric mouse SAA3) (38, 39). Multiple oligomeric species of SAA1 and SAA2 have been reported, with one study showing initial formation of a soluble SAA octamer followed by conversion to a hexameric species over time (40).

While the tetrameric structure of mouse SAA3 has been suggested to transport retinol via a hydrophobic pocket formed by four copies of helix 3, no clear functionality has been attributed to the hexameric form of SAA1 (40). The apex of the SAA1 hexamer contains a cluster of positively charged residues proposed to play a role in SAA interactions with glycosaminoglycans (GAGs) and with HDL, although most molecular details of GAG or lipid binding have been determined using truncated forms of SAA (41, 42). Within the SAA1 hexamer structure, the C-terminal region forms a loop that makes a wide range of salt bridges and hydrophobic contacts with the four helix bundle, in contrast to the proposed disorder of this key receptor-binding domain (38).

It has been confirmed by NMR, SAXS, and fluorescence spectroscopy that ApoA-I and membrane scaffold protein (MSP), a derivative of ApoA-I, bind to discoidal HDL or HDL-like particles by wrapping around the perimeter as a helical-belt, interacting with the exposed hydrophobic bilayer core (43–45). To date however, there have been no experimental studies demonstrating the structure of lipid-bound SAA. Based on the compact SAA monomer (a cone-shaped four-helix bundle) that makes up the oligomers in reported crystal structures, it has been predicted that SAA binds to the surface of lipid bilayers in this monomeric form, with the hydrophobic faces of helices 1 and 3 directly contacting the lipid. In this model, the more disordered C-terminal region faces away from the membrane, to mediate receptor interactions (46, 47). The binding of a compact monomer of SAA to the surface of an HDL particle does not account for how SAA is capable of displacing ApoA-I or for the more discoidal shape reported for SAA containing HDL, with the latter suggesting an ApoA-I like mechanism of SAA interaction with HDL (48).

Understanding the structural properties and interactions of both lipid-free and lipid-bound SAA are important for understanding their roles in health and disease. In fact, the biological role of lipid-free SAA remains controversial despite surface plasmon resonance studies of SAA-HDL interactions suggesting the presence of appreciable amounts of free SAA during the acute phase response (36, 49). Here, we report structural models for both a previously uncharacterized lipid-free SAA octamer and for SAA-lipid nanodiscs scaffolded by tetrameric SAA. These models are supported by a combination of biochemical crosslinking, small angle X-ray scattering (SAXS), and analysis of transmission electron microscopy (TEM) images obtained for both states of SAA. Together, these data describe an open ring-like SAA structure that is capable of scaffolding discoidal lipid particles in a manner similar to ApoA1. Thus, we provide new insight into SAA-lipid interactions that may play important roles in both SAA-HDL interactions during the inflammatory response and suggest a mechanism through which SAA might act to sequester and remove lipids from sites of tissue damage.

## Experimental procedures

### Recombinant expression and purification of serum amyloid A

Recombinant human SAA2.1 and SAA1.1 (11.7 kDa) were expressed in BL21 (DE3) E. coli. In each case, N-terminally His_6_-tagged SAA was purified from inclusion bodies using nickel-affinity chromatography in denaturing conditions (6 M GuHCl, 250 mM NaCl, 50 mM sodium phosphate pH 8.0) at room temperature. Purified SAA was dialyzed into a refolding buffer (400 mM L-Arginine, 1.1 M GuHCl, 21 mM NaCl, 0.88 mM KCl, 1 mM EDTA, 55 mM Tris pH 8.2) overnight at 4°C. Refolded protein was further dialyzed into low salt phosphate buffer (1 M urea, 21 mM NaCl, 20mM sodium phosphate pH 7.5) overnight at 4°C. His-tag removal was achieved through cleavage with thrombin for 5 hrs at room temperature. Thrombin was removed by running the cleaved sample on a benzamidine-sepharose column, residual thrombin activity was stopped using PMSF and SAA was dialyzed against 20 mM Tris (pH 8.0) with 150 mM NaCl, concentrated, and further purified using size exclusion chromatography (Superdex 200 10/300 GL). The purity of the SAA containing fractions was checked with SDS-PAGE.

### Formation of SAA-lipid nanodiscs

Dimyristoyl phosphatidylcholine (DMPC) and dimyristoyl phosphatidylserine (DMPS) in chloroform were dried using nitrogen gas and residual solvent was evaporated under vacuum. Lipids were suspended in water, subjected to several freeze-thaw cycles, lyophilized overnight, and then resuspended in 20 mM Tris buffer (pH 8.0) to form multilamellar vesicles (MLVs) composed of 80:20 mol ratio of DMPS:DMPC (PS:PC). MLVs were bath sonicated and extruded using 0.2 μm filtration membrane to form small unilamellar vesicles (SUVs), and stored at 37°C. The extruded SUVs were incubated with freshly purified SAA2.1 at mass ratio of 1:1 protein:lipid for 3 hours at 26°C to form lipid-bound SAA2.1 nanodiscs. Both SAA2.1-DMPC and SAA1.1-DMPC nanodiscs were used for visualization with electron microscopy while only the monodisperse SAA2.1-PC:PS particles were used for small-angle X-ray scattering studies.

### Circular dichroism (CD) spectroscopy

Far-UV CD spectra were recorded using a Jasco J-810 spectropolarimeter. SAA2.1 or SAA1.1 samples were prepared (0.2 mg/ml protein in 20 mM Tris, pH 8.0) and loaded in a quartz cuvette (0.1 cm path length). Spectra were collected at 4 °C and represent an average of three scans from 190-260 nm.

### Blue native and clear native gel electrophoresis

For blue-native (BN)-page, protein samples (0.2 mg/mL) were diluted in a 2x gel loading buffer (500 mM 6-aminocaproic acid, 150 mM bis tris-HCl pH 7.0, 100 mM NaCl, 20% glycerol, 1% w/v Coomassie blue G-250) and loaded on a 15% polyacrylamide bis-tris gel. The cathode buffer (0.2% w/v Coomassie G250, 500 mM tricine, 150 mM bis-tris pH 7.0) and anode buffer (500 mM bis-tris pH 7.0) for running BN-page were made from 10x stock solutions. The gels were run at 160 V for 70 min and then stained with Coomassie dye. Clear native gels were run similarly but in the absence of Coomassie blue G-250 in the loading and running buffers.

### Glutaraldehyde crosslinking followed by SDS-PAGE

Lipid-free and lipid-bound SAA (0.3 mg/mL) were mixed with 0.5 μL of 2.5% glutaraldehyde at different times (37 °C). The crosslinking was terminated by the addition of 1 μL of 1 M Tris solution followed by dilution in a 2x gel loading buffer (500 mM 6-aminocaproic acid, 150 mM bis tris-HCl pH 7.0, 100 mM NaCl, 20% glycerol, 1% w/v Coomassie blue G-250). The final samples were loaded on a 15% polyacrylamide tris gel. The gel was run at 160 V for 70 min and then silver stained.

### Trypsin digestion followed by SDS-PAGE

Lipid-free and lipid-bound SAA (0.3 mg/mL) were mixed with Trypsin (1:1000 trypsin:protein) at different times (37°C), and the digestion was terminated by the addition of the 2x gel loading buffer (500 mM 6-aminocaproic acid, 150 mM bis tris-HCl pH 7.0, 100 mM NaCl, 20% glycerol, 1% w/v Coomassie blue G-250). The final samples were loaded on a 15% polyacrylamide tris gel. The gel was run at 160 V for 70 min and stained with Coomassie dye.

### Size-exclusion chromatography with multi-angle light scattering (SEC-MALS)

SAA samples and bovine serum albumin (BSA) controls were separated at 4°C on a Superdex 200 10/300 column at a flow rate of 0.5 ml/min. Refractive index measurements were obtained using an Optilab T-rEX, and multi-angle light scattering was obtained using a miniDAWN TREOS WYATT (Wyatt Technology, Santa Barbara, CA). SEC-MALS data were processed and analyzed using ASTRA (version 7.1.3).

### Small-angle X-ray scattering (SAXS)

An Anton Paar SAXSpace instrument with SAXSDrive software was used to collect SAXS scattering curves for both the lipid-free and lipid-bound forms of SAA2.1 in 20 mM Tris buffer (pH 8.0). An exposure time of 30 min/frame and a total of 48 frames were used. The sample to detector distance was 317.0598 mm. The data for lipid-free SAA2.1 were recorded at 10°C while lipid-bound SAA2.1 data were recorded at 27°C (above the 25 °C gel-to-liquid phase transition temperature of DMPC). The one-dimensional data contain the scattered intensity as a function of the scattering vector magnitude *q* = (4π/λ)sinθ or *S* = (2π/λ)sinθ, where 2θ is the scattering angle and λ is the X-ray wavelength. The zero position for each scattering curve was calibrated using SAXSTreat. Further processing such as buffer subtraction and desmearing of the line collimation data was done using SAXSquant.

Analysis of the processed SAXS data was done using tools from the ATSAS2.6 package (EMBL, Hamburg, Germany). PRIMUS was used to perform Guinier analysis, and to visualize Kratky plots (50). GNOM was used to convert data from reciprocal space to real space by Fourier transformation and generate pair-distance distribution functions (51). The resulting GNOM output file was used to generate a low-resolution molecular envelope using GASBOR (52). Consequently, DAMAVER was used to build a final averaged model (53). CRYSOL was used to both generate scattering curves from the PDB structures and to compare them to experimental scattering curves (54).

### Transmission electron microscopy

Lipid-free SAA2.1 (purified by size-exclusion chromatography) was adsorbed to glow-discharged carbon-coated copper grids at 4°C. Grids were subsequently washed with deionized water and stained with freshly prepared 0.75% uranyl formate. Lipid-bound SAA2.1 samples were prepared in a similar manner but were stained with 2% uranyl acetate. Electron micrographs for all specimens were obtained with a FEI Technai 20 electron microscope operated at 120 kV. For single particle analysis of the negative stain TEM data, 1005 SAA2.1 particles from 92 images, and 1303 lipid bound SAA2.1 particles from 105 images were used. Relion (version 2) 2D classification was used, resulting in 10 class averages for each form of SAA (55).

## Results

### Oligomeric state of human SAA in solution

Serum amyloid A has an inherent tendency to aggregate during purification in its native form, giving rise to multiple possible oligomeric states and confounding downstream analysis (40, 56). To avoid aggregation, we purified SAA (both the human SAA1.1 and SAA2.1 isoforms, with sequences shown in Figure S1a under denaturing conditions, followed by refolding, resulting in a stable, soluble state. Soluble SAA was then mixed with DMPC or DMPS SUVs to form SAA-lipid nanodiscs. The secondary structure of both the lipid-free and the lipid-bound forms of SAA were analyzed using CD spectroscopy. The spectra shown in Figure 1a are consistent with predominantly α-helical structures for both the lipid-free and lipid-bound forms of SAA, consistent with the previously reported high-resolution structures of SAA (38, 39). No difference in CD spectrum was observed between SAA1.1 and SAA2.1 (Figure S1b), suggesting that the small sequence differences between these major isoforms do not have a significant impact on SAA structure.

**Figure 1.**
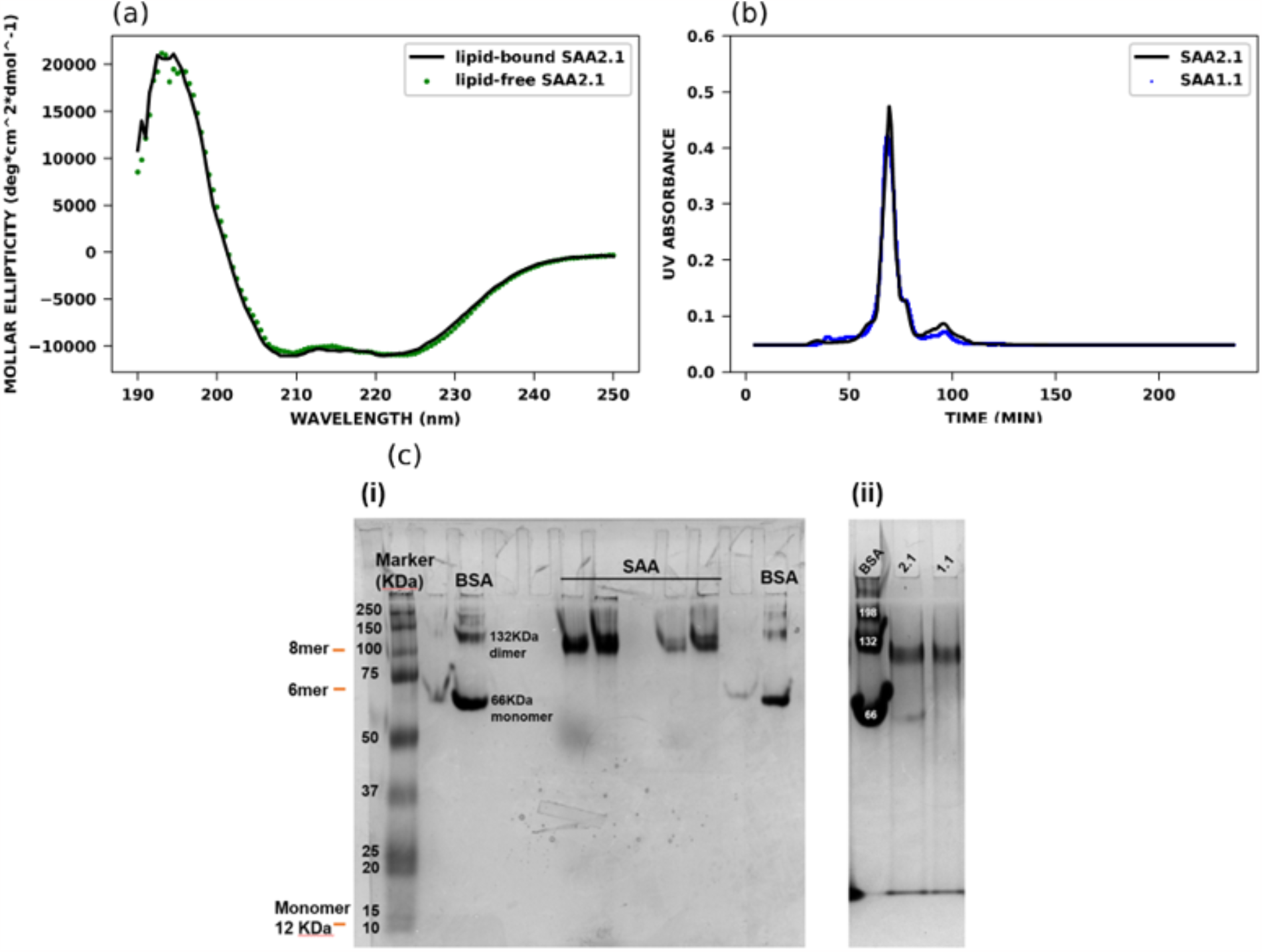
Secondary structure and oligomeric state of lipid-free and lipid-bound SAA. (A) Circular dichroism spectra of 20μM lipid-free and lipid-bound SAA in 20mM Tris (pH 8) at 25 °C. The spectrum is an average of 3 scans. (B) Size exclusion chromatography-multi angle light scattering (SEC-MALS) analysis of lipid-free SAA1.1 and SAA2.1 in 20mM Tris, 150 mM NaCl (pH 8). (C) (i) Clear native-PAGE (15%) of SAA2.1 and (ii) blue native-PAGE (15%) analysis of SAA2.1 and SAA1.1 in 20mM Tris (pH 8).

Although the high degree of α-helical content observed here is consistent among structural studies of SAA, there are varying reports on the oligomeric state of SAA, including tetrameric, hexameric, and octameric states (38–40, 57). This has been previously attributed to differences in isoform used, presence/absence of a purification tag, and variations in solution conditions. To determine the oligomeric state of our recombinant SAA2.1 and SAA1.1 proteins, we performed clear native-PAGE, blue native-PAGE, and SEC-MALS analysis on the lipid-free form of both isoforms (Figure 1b-c). Figure 1c(i-ii) shows that the migration of refolded SAA in both clear and blue native-PAGE is consistent with a predominant oligomeric state that is larger than would be expected for with tetrameric (48 kDa) or hexameric (72 kDa) SAA. Comparison with the migration of molecular weight standards and bovine serum albumin (BSA) suggests a molecular weight close to 100 kDa for our SAA oligomers, with a minor band at approximately ½ of this mass. Note that unlike the SAA 2.2 octamers reported by Wang et al., our SAA 1.1 and 2.1 octamers remained stable over days-weeks and did not convert to hexamers (40).

SEC-MALS was used to more accurately determine the molecular weights of SAA2.1 and SAA1.1. This technique uses light scattering at three different angles to extrapolate the scattering at zero angle, which is proportional to the molecular weight and concentration of protein. The concentration of the separated species is measured through UV absorption and refractive index, allowing mass and oligomeric state to be unambiguously determined if the extinction coefficient for the protein and dn/dc (difference in refractive index between sample and solvent) are known. Figure 1b shows that both SAA2.1 and SAA1.1 have the same retention time on size exclusion, with a MALS-determined molecular weight of (96 ± 1.13%) kDa, which is consistent with SAA existing as an octamer (monomer mass = 12 kDa). There is also a less prominent peak to the right of the 96 kDa peak, corresponding to a 48 kDa tetramer and suggestive of lipid-free SAA existing as a dimer of tetramers. The faint band at approximately 50 kDa in our native-PAGE data is consistent with this analysis, providing further support for this model. Since both SAA1.1 and SAA2.1 exhibited the same secondary structure and oligomeric behaviour, subsequent structural studies focused only on SAA2.1.

### Structure of lipid-free SAA in solution

Serum amyloid A plays an important role in both innate immunity and lipid transport, and has been shown to interact with a variety of immune receptors through its C-terminal region (46, 58, 59). Thus, we fed the sequence of SAA2.1 into a disorder predictor neural network (MetaPrDOS) to determine whether the C-terminal region contains any disorder that might account for this promiscuity(60). Figure S2a shows that indeed there is higher disorder predicted at the C-terminal region (amino acids 76-106), suggesting that residues 1-76 are largely helical to account for our CD results. This result is also consistent with the higher crystallographic B-factors observed for this region by Lu et al. (2014), and with the region that is cleaved through proteolysis during AA amyloidosis to produce fibrils containing only amino acids 1-76 (38).

SAXS was used to define the overall structure of lipid-free SAA octamers in solution. Figure 2a shows the scattering curve for the lipid-free SAA2.1. The presence of a well-developed minimum at the scattering vector q of 1.2 nm^-1^ indicates that the sample is monodisperse and that the octameric complex is not linear in shape or intrinsically disordered. The SAA octamer has a radius of gyration (R_g_) of 3.2 nm as determined from Guinier analysis (Fig. 2b) and based on the pair-distance distribution function (PDDF), has a maximum dimension of approximately 11.7 nm (Fig. 2c). The PDDF is reminiscent of a Gaussian distribution but has an extended tail that suggests an elongated spherical shape to SAA. Moreover, the highly structured nature of SAA octamers was further confirmed through Kratky analysis (Fig. 2d). While folded proteins give rise to a bell-shaped curve and a return to zero, the Kratky plot of a disordered protein does not return to baseline. Thus, the Kratky plot for SAA indicates a predominantly ordered protein, with the potential for some disordered segments, as evidenced by the slightly elevated values at high q, in agreement with our CD and disorder prediction data. The scattering curve obtained for lipid-free SAA was compared to back-calculated SAXS data for the reported PDB structures of tetrameric mouse SAA3 (PDB: 4Q5G, Fig. S3a) and hexameric human SAA1.1 (PDB: 4IP9, Fig. S3c). Neither are a good match for our experimental data, indicating that our octamers represent a previously uncharacterised structure for lipid-free SAA (38, 39).

**Figure 2.**
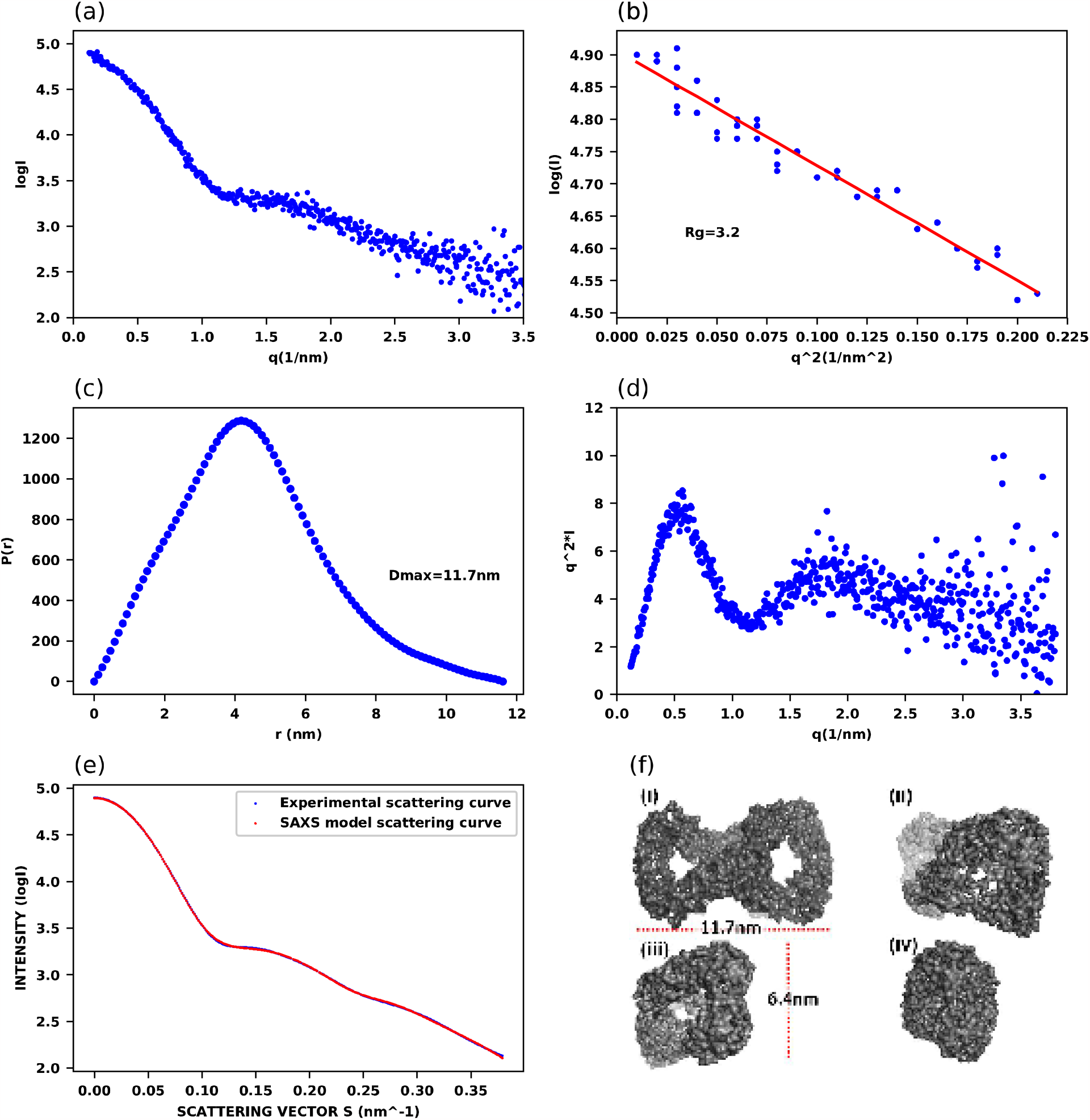
Small angle X-ray scattering of lipid-free hSAA2.1. (A) SAXS scattering curve of SAA2.1 in 20mM Tris (pH 8). (B) Guinier approximation of SAA2.1 scattering data (C) Pair-distance distribution of SAA2.1 with maximum distance of 11.7nm. (D) Kratky plot of SAA2.1 scattering data. In-house Anton Paar SAXSpace instrument was used to collect the scattering curve shown in Fig.3a. (E) Fit of the ab-initio reconstructed model to the experimental scattering curve and the (F) (i-iv)) Different views of the ab-initio reconstructed model using GASBOR.

Ab initio modeling was used to generate a low-resolution molecular envelope of octameric SAA from the SAXS data, using an imposed P2 symmetry based on our data suggesting that octameric SAA may be a dimer of tetramers. Four different views of the resulting 3D model are shown in Figure 2f (i-iv), and a scattering curve generated from this model shows a high alignment with the experimental scattering curve obtained for SAA (Figure 2e, χ^2^=1.4). The overall structure of the SAA octamer resembles the figure “8” or infinity sign, which is very similar to the twisted ring-like structure of lipid-free ApoA-I tetramers reported by Borhani et al., although the latter has a somewhat more open conformation compared to SAA (61). Due to this difference, the simulated SAXS scattering curves of ApoA1 is not a good match to the SAA data (Figure S3b) highlighting that these proteins form distinctly different structures. Interestingly, in both cases, the structures have a similar ‘belt’ thickness of approximately 2 nm. The tetrameric structure of ApoA-I contains 4 alpha-helices within this 2 nm thick region, and is approximately twice as long as SAA, further supporting an octameric SAA species (a 199-residue fragment containing amino acids 44-243 of Apo A-I was used for the tetramer structure (PDB: 1AV1), relative to the 106 amino acid SAA used here) (61). The close contact between the four alpha helices can also explain the observation of a tetramer in our native PAGE and glutaraldehyde crosslinking experiments.

To provide additional confirmation of the ab initio model obtained from our SAXS data, we used single particle analysis of negative stain TEM images. Figure 3a shows a representative micrograph of the SAA octamers, with an expanded image of a single particle shown in the inset. Class averages of 1005 SAA particles are shown along with a set of manually selected views from the 2-fold symmetric SAA SAXS model (Fig. 3b-c). There is good visual agreement between the different classes of SAA particle obtained from TEM and various rotated views of the SAXS model, increasing confidence in our low-resolution twisted-belt structure of a soluble SAA octamer.

**Figure 3.**
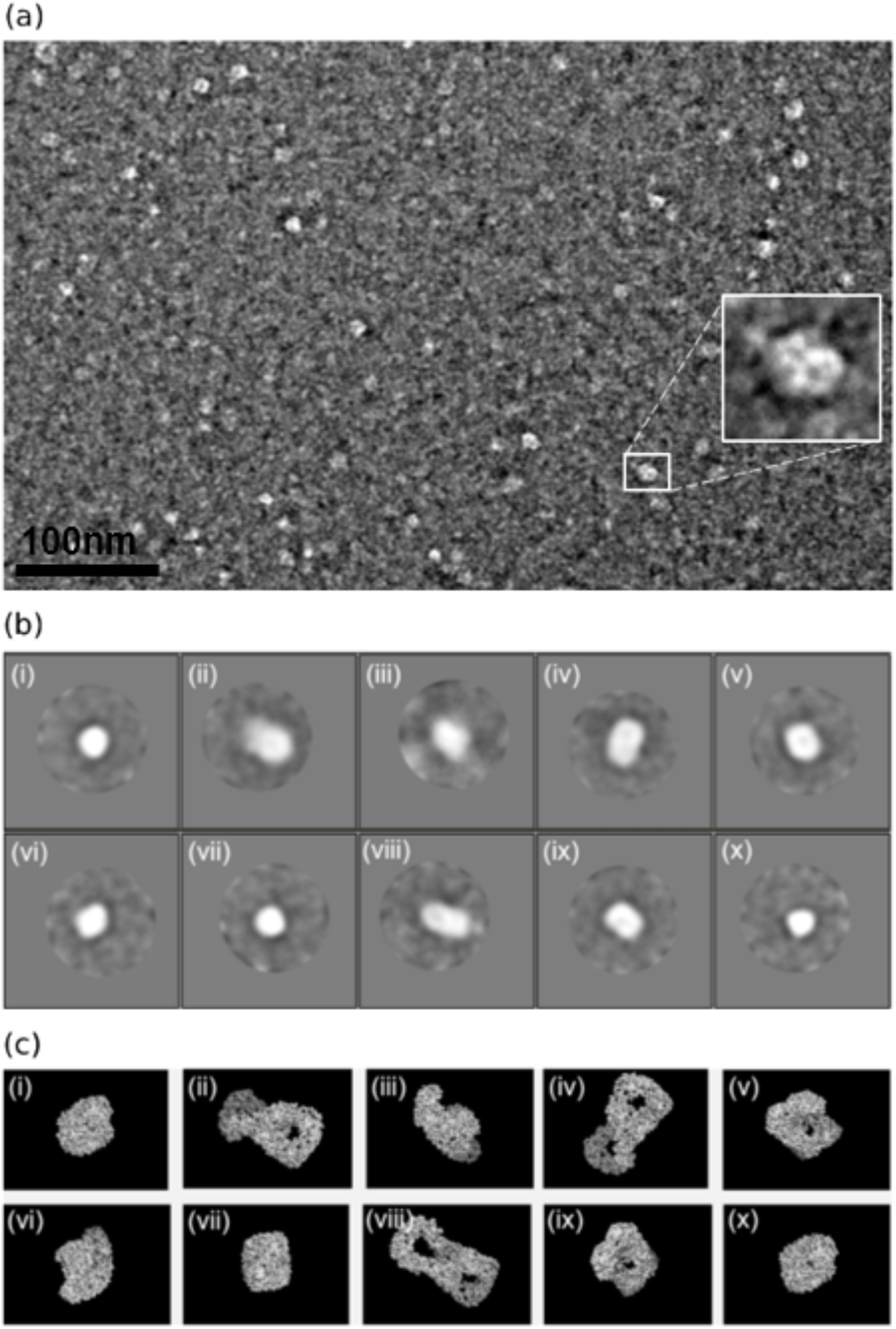
Transmission electron microscopy of lipid-free hSAA2.1. (A) Representative micrograph of hSAA2.1 stained with 0.75% (w/v) uranyl formate. (B) 2D class averages of hSAA2.1 from 1005 selected particles and the (C) corresponding views of the ab-initio model of hSAA2.1 generated from SAXS data.

### The structure of SAA-lipid nanodiscs

When added to SUVs composed of either DMPC or a mixture of 80:20 mol ratio DMPS:DMPC, the octameric lipid-free SAA remodels the lipid from 0.1-0.4 micrometre-sized liposomes into small nanodiscs with regular shape and size (Figure 4a, Figure S4). Note that while the dimensions and morphology of individual SAA-lipid nanodiscs are the same for both DMPC and DMPS:DMPC containing particles, those formed in the absence of anionic lipids exhibited a strong tendency to assemble into stacked arrays, while DMPS containing nanodiscs were monodisperse in solution. In both cases, the SAA-lipid nanodiscs closely resemble those formed through incubation of either Apo A-I or MSP with lipid (62, 63). A similar negative-stain TEM approach was used to analyze the SAA nanodiscs as described above for the lipid-free protein. Figure 4b shows 10 class averages of lipid-bound SAA nanodiscs from a total of 1303 picked particles. Only unstacked nanodiscs were used in this analysis. Note that these class averages lack the level of structural detail observed for SAA octamers in the absence of lipid, likely reflecting challenges in using negatively stained protein-lipid particles as well as potential conformational dynamics inherent in this system. The average diameter of a lipid-bound SAA nanodiscs is approximately 12.3 (± 1.9) nm (105 discs measured using ImageJ, NIH), with a particle thickness of 2.5 (± 0.34) nm. Based on the well-established modes of lipid binding by Apo A-I and MSP, and the twisted ring-like structure of octameric SAA, we hypothesized that SAA likely binds and remodels lipid in a similar manner by wrapping around lipid in a helical-belt conformation.

**Figure 4.**
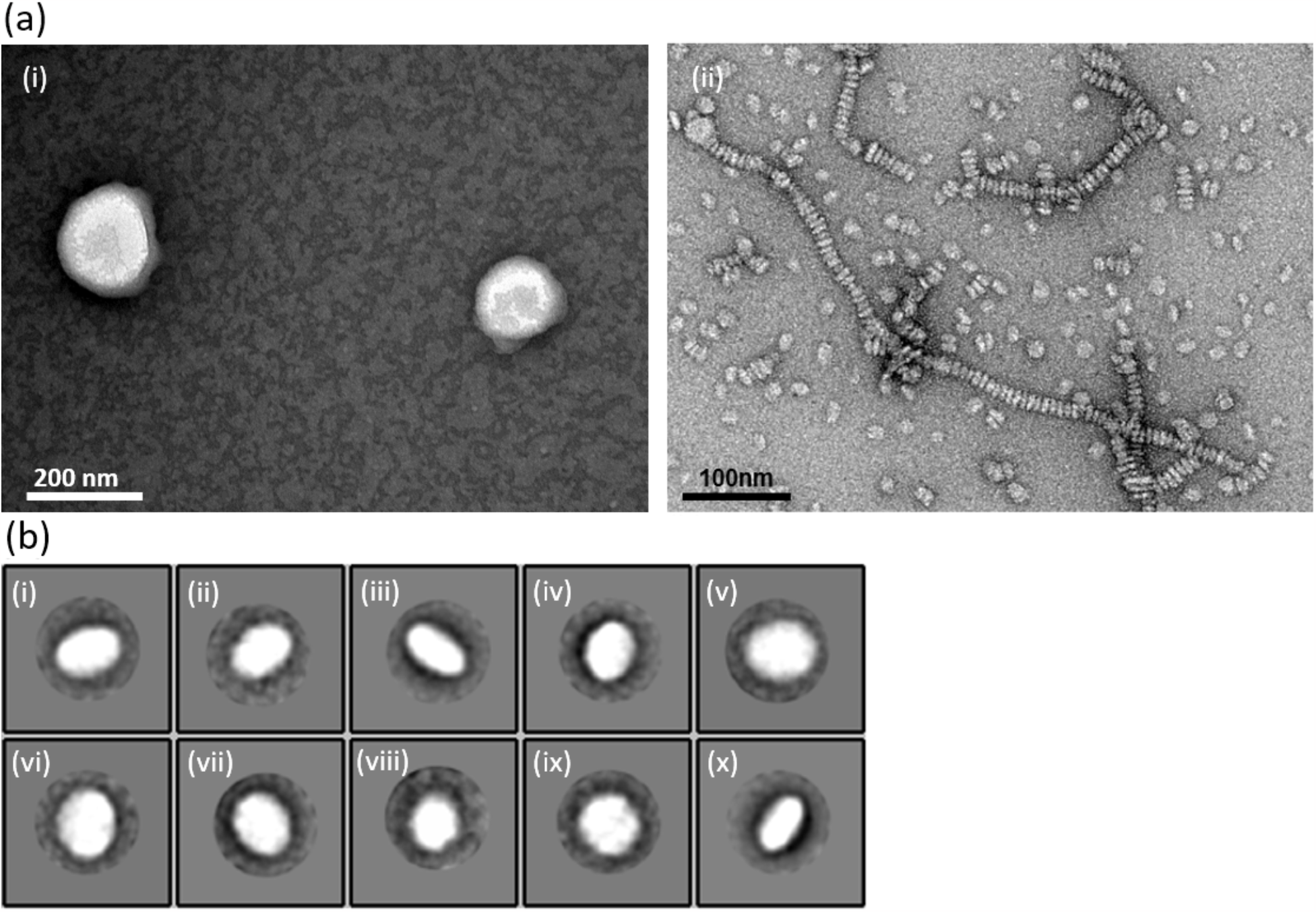
Transmission electron microscopy of lipid-bound hSAA2.1. (A) Representative micrograph of a (i) liposome and (ii) lipid-bound SAA stained with 2% (w/v) uranyl acetate. Discoidal HDL-like nanoparticles shown were formed after incubation of 1:1 DMPC-to-SAA2.1 (w/w) after 2 hrs at 26°C. (B) Class averages of lipid-bound hSAA2.1 nanoparticles from a total of 1303 selected particles.

To test this hypothesis, we performed SAXS experiments on the SAA nanodiscs in solution, preparing them with negatively charged DMPS lipid to avoid the stacking observed for DMPC nanodiscs in Figure 4. Figure 5a shows the scattering curve obtained for the SAA-DMPS nanodiscs, the monodispersity of which were confirmed using TEM (Figure S4a) and through Guinier analysis (Figure 5b) in which a linear fit indicates a lack of aggregation or repulsion in the system. From the Guinier analysis, the R_g_ of the SAA nanodiscs was determined to be 3.96 nm, which is larger than that of lipid-free SAA, suggesting that SAA in the lipid-bound state has a more open conformation, suggesting some rearrangement is required during lipid scaffolding. The PDDF of SAA nanodiscs shows a second peak (Fig. 5c), which is indicative of a distinct two-component morphology. In the case of aggregation or two-domain particles, the second peak will be smaller due to the large proportion of close distances within each particle or domain and only a few long distances connecting the two subunits. The PDDF of lipid-bound SAA, however, shows a larger second peak, which is generally indicative of a core-shell particle. In addition to representing all interatomic distances in a particle, the PDDF is weighed by density, such that these data suggesting that the SAA nanodiscs are discoidal particles with higher density at longer distances, arising from a peripheral belt of protein, and a low-density lipid core giving rise to the shorter distances. The PDDF analysis, therefore, supports the idea that SAA scaffolds lipid nanodiscs, and potentially discoidal HDL, by uniformly wrapping around it in a helical-belt conformation similar to ApoA1 or MSP. Moreover, The Kratky plot of lipid-bound SAA nanodiscs shows a curve characteristic of a folded protein. This in addition to CD data indicate that SAA maintains its folded alpha helical secondary structure while bound to lipid (Fig. 5d).

**Figure 5.**
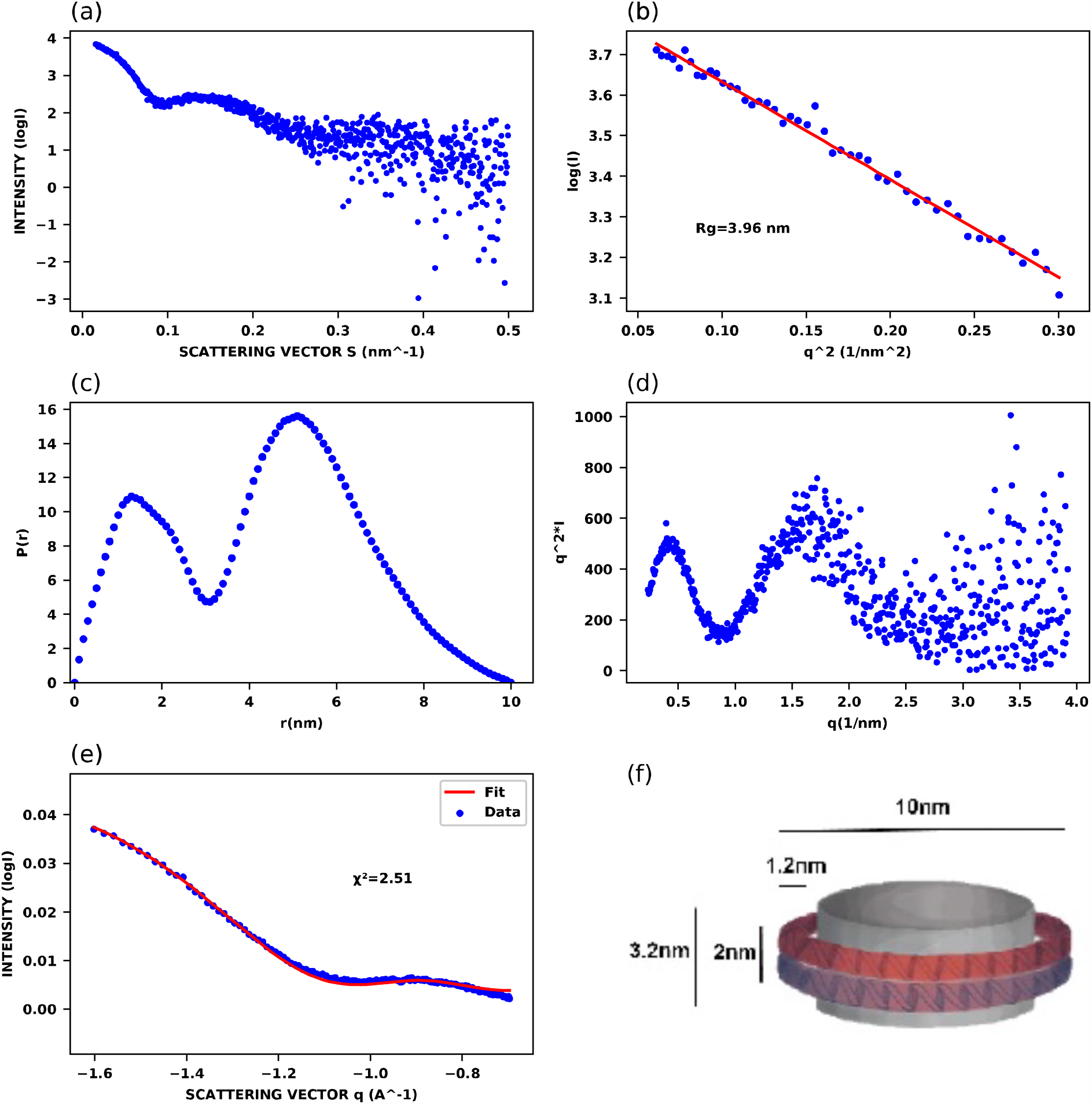
Small angle X-ray scattering of lipid-bound hSAA2.1. (A) SAXS scattering curve of 4mg/mL lipid-bound SAA in 20mM Tris (pH 8). Nanoparticles were formed using a mixture of DMPS:DMPC (0.8:0.2) to reduce stacking and improve monodispersity. (B) Guinier approximation of lipid bound-SAA (points10-49). (C) Pair-distance distribution of lipid-bound SAA and (D) Kratky plot of lipid-bound SAA scattering data. (E) Comparison of the scattering curve of lipid bound SAA2.1 (lipid = DMPS:DMPC 80:20) to the nanodisc model of the *WillItFit* software. (F) Schematic of lipid-bound SAA depicting the relative dimensions of the protein belt and the lipid bilayer determined using SAXS and *WillItFit*.

We used CRYSOL to compare our experimental scattering curve of SAA-lipid nanodiscs to that calculated for nanodiscs formed by lipid-bound MSP. Figure S3d shows that the two scattering curves are very similar in shape (χ^2^ value of 0.407) and both share a dip at approximately q = 0.1 nm^-1^ and a peak ranging from 0.14 - 0.2 nm^-1^. The similarity between the two scattering curves provides further support towards SAA binding to discoidal HDL-like particles in a similar manner to both MSP and Apo A-I. The relatively small difference between the two scattering curves may be due to minor structural differences caused by variations in the number or fluidity of the lipid component or the dimensions of the protein belt. Alternatively, the two proteins as well as due to the experiments being done at slightly different temperatures of 27 °C (SAA) vs 30 °C (MSP).

### Oligomeric state of SAA-lipid nanodiscs

Ab initio modelling of lipid-bound SAA based on the SAXS data was not feasible as it is a two-component system consisting of lipid and protein with two different densities. Therefore, to characterize the structural features of lipid-bound SAA we made use of the software *WillItFit* (64). The parameters defining the resulting nanodisc structure are provided in Table S1, and include volume, the number of lipids (60), height of the protein belt (2nm), and height of the lipid bilayer bilayer (3.2nm) as calculated using *WillItFit*. A schematic of the SAA-lipid nanodisc model is shown in Figure 5f. Validation of this model was carried out by comparing the experimental SAXS data for lipid-bound SAA with that calculated for the nanodisc model. As shown in Figure 5e, we found that lipid-bound SAA fits to our predicted nanodisc model with a χ^2^ value of 2.5 (Figure 5e) supporting the TEM and SAXS data and providing additional evidence that the mechanism of lipid binding by SAA is similar to that of Apo A-I and MSP. Slight discrepancies between the experimental scattering data and that calculated for the *WillItFit* model likely arise from deviations from ideal geometry in the experimental particles, and from assumption that the protein forms an ideal, fully lipid-bound helical belt.

Based on the volume of the belt divided by the volume of one SAA monomer, the expected number of SAA proteins is calculated to be 3 fully bound SAA monomers or 4 partially bound monomers. To better define the number of SAA molecules per nanodiscs, we performed both crosslinking experiments (Figure S5) and trypsin digestion followed by SDS PAGE (Figure S6) on lipid bound SAA. Figure S5 compares glutaraldehyde crosslinking results for lipid free SAA, DMPC-bound SAA, and DMPC:DMPS-bound SAA. Lipid-free SAA initially presents as a ladder from monomer to octamer molecular weight, but is rapidly aggregated by the crosslinking reaction. A similar result is observed for the SAA-DMPC nanodiscs, with monomer, dimer, trimer and tetramer showing as the predominant species, along with some aggregate and higher molecular weight species that likely result from inter-disc crosslinking as a consequence of stacking. This aggregation is absent for SAA-PC:PS nanodiscs, which show a slower progression of crosslinking and no species larger than tetramer (monomer, dimer and trimer are the strongest bands). These data match the modeling results, and suggest that three or four molecules of SAA are present in each nanodisc. Note that the similarity of the SAA nanodiscs with those formed by APOA-1, which can form extended helical structure twice the length of those formed by SAA might indicate a requirement for four SAA molecules to scaffold lipid nanodiscs.

To determine if the discrepancy between 3 and 4 SAA monomers per nanodisc reflected in the *WillItFit* results was due to the presence of disordered or non-lipid bound regions of SAA, trypsin digestion was performed to look for exposed or flexible protein segments. As shown in Figure S6, the lipid-free form of SAA is completely digested after 5 minutes while SAA bound to DMPC is significantly stabilized, by the protein-lipid interactions and requires 10-30 minutes to become fully digested by trypsin. In both lipid-bound SAA samples a second species is quickly produced that is approximately 1 kDa smaller than the full-length monomer. This species represents the more stable, and more likely lipid-bound, core of SAA in the nanodiscs. Interestingly, this species is also present in the lipid-free SAA at shorter incubation times, although it is subsequently degraded by trypsin. This strongly suggests that the labile segment of SAA is a portion of the C-terminal region (amino acids 76-107) that is predicted to exhibit higher disorder, and which has been implicated in SAA binding to immune receptors, glycosaminoglycans, among other intermolecular interactions (42, 58, 59). Together, our data support dissociation of a soluble SAA octamer into two tetramers upon interaction with lipid vesicles, each of which can scaffold an HDL-like nanodisc. These SAA tetramers likely form a helical belt around the periphery of the particle, with the C-terminal region of each SAA monomer remaining unbound.

## Discussion

SAA has been implicated in several of processes that take place during the inflammatory response, stimulating production of pro-inflammatory cytokines and matrix metalloproteinases as well as promoting immune cell migration into damaged tissues. Additional proposed roles for SAA include altered reverse cholesterol transport properties of SAA-containing HDL, opsonization of bacteria, and clearance of lipids from damaged tissue (27, 65–67). The ability of a relatively small protein to exhibit such diverse activity may be facilitated by the presence of two major isoforms (SAA1 and SAA2) and by different activity of lipid-free and lipid-bound forms of the protein. While it has been estimated that up to 95% of SAA in circulation during the acute phase response is associated with lipoproteins, less is known about the state of SAA produced during long-term inflammation or in non-hepatic tissues (68). Just as multiple oligomeric states have been observed for lipid-free SAA, multiple lipid-bound forms may also exist in vivo, allowing functional interactions of SAA with HDL, other lipoproteins, or free phospholipids.

Prior structural studies of SAA have identified a stable, compact hexamer in which each monomer is a helical bundle with a slightly curved conical shape. These hexamers are distinct from the open, twisted ring octamers revealed by our data. These octamers, which are reminiscent of ApoA-I, may reflect a less stable state of the protein, although we did not observe evidence of instability or conformational change over time in our samples (40). Such metastability or change in oligomeric state may be highly dependent on solution conditions. Interestingly, while no shift to hexamer or to high order aggregates was observed in our samples, the open structure of the SAA octamers suggests a more flexible structure than the hexamer, with more potential for rearrangement of helical contacts. Similarly, there was no isoform dependence, with both SAA1.1 and SAA2.1 forming octamers in the absence of lipids. The ab initio model of lipid-free SAA2.1 built using our SAXS data shows important similarities to the structure of lipid-free ApoA-I, having an open structure formed by an approximately 2nm thick helical assembly. With a strong body of evidence indicating that SAA displaces at least some fraction of the ApoA-I from HDL during the acute phase response, it is likely that this structural similarity has functional implications, potentially priming SAA to make extended helical contacts with lipid membranes (69).

A functional role for this open SAA octamer structure is supported by our modeling of SAA-lipid nanodiscs. The discoidal HDL-like particles formed here by SAA and DMPC or DMPS are similar in overall morphology to those previously reported by Takase et al., who also observed that lipid-free and lipid-bound SAA had similar helical content (33). Frame et al. also described discoidal nanodiscs formed by SAA and POPC and, based on the overall dimensions of their particles and estimates of the lipid:protein ratio, proposed a model in which 12 molecules of SAA were required to scaffold 72 lipid molecules (70). Based on our TEM, SAXS and crosslinking data, we have determined that a tetramer of SAA is sufficient to form nanodiscs containing ~60 molecules of DMPC. Further, the dimensions of our SAA-lipid particle can be related to the lipid free octamer structure, such that an opening of the figure-eight conformation to form a ring, followed by dissociation into tetramers would be sufficient to allow formation of nanodiscs. We hypothesize that similar to ApoA-I and MSP nanodiscs, two stacked α-helices are required to wrap around the hydrophobic circumference of these particles, such that an appropriately arranged tetramer of SAA is able to replace a dimer of the longer ApoA-I protein (62, 63, 71).

Other models of SAA-lipid interaction based on the hexamer crystal structure have been proposed based on identification of a curved hydrophobic surface that would be exposed if the hexamer dissociated into cone-shaped monomers (46). In this model, the curvature of the monomer is proposed to better match that of HDL particles, with lower affinity for less curved LDL or VLDL. Our current observations provide experimental support for an alternative model for SAA-lipid interactions, which might allow for more flexibility in lipoprotein binding. For instance, it has been found recently that in patients with chronic-inflammation, LDL-bound SAA is present (72, 73). Our lipid-belt model also suggests a mechanism by which SAA could displace ApoA-I from HDL, consistent with observations of reduced ApoA-I content in SAA-bound HDL particles (74, 75). The ability of octameric SAA to scaffold lipids in this way also suggests a likely mechanism for SAA-mediated lipid clearance during inflammation, which would require significant structural rearrangement of the hexameric lipid-free form of the protein (15, 70). Given that lipid-free SAA is capable of adopting multiple structures, more than one type of lipid-SAA interaction may also be adopted as a means to facilitate the distinct functions of SAA in inflammation, innate immunity and lipid clearance.

In conclusion, we have reported new structural models for lipid-free and lipid-bound human SAA, based on microscopy, scattering and biochemical data. The lipid free octamer closely resembles solution structures of ApoA-1 and is capable of dissociating into tetramers in the presence of DMPC or DMPS, allowing it to form a helical belt to stabilize protein-lipid nanodiscs. The lipid-free model presented here represents a new soluble form of SAA and demonstrates the ability of this protein to adopt multiple stable structures in the absence of lipids, which has potential implications for the diverse functionality attributed to this small protein. Similarly, we present the first experimentally derived model for lipid-bound SAA, showing how this protein can form a helical belt scaffold for lipid-protein particles that resemble very small discoidal HDL, similar to the nanodiscs that are scaffolded in a similar manner by ApoA-I or MSP (62, 63). This model provides a new perspective on SAA-lipid interactions and suggests new mechanisms for some of the reported activities of this protein, including the displacement of ApoA-I from HDL particles and clearance of lipids from sites of tissue damage.

## Supporting information

This article contains supporting information.

## Acknowledgements

The authors acknowledge the use of The Hospital for Sick Children’s Structural & Biophysical Core Facility and Nanoscale Biomedical Imaging Facility.

## Funding and additional information

This work was supported by Natural Sciences and Engineering Research Council of Canada Discovery Grant RGPIN 06146 (to SS). AN was supported by a SickKids Restracomp scholarship.

