## Supporting Information for "Structural studies of a serum amyloid A octamer that is primed to scaffold lipid nanodiscs"

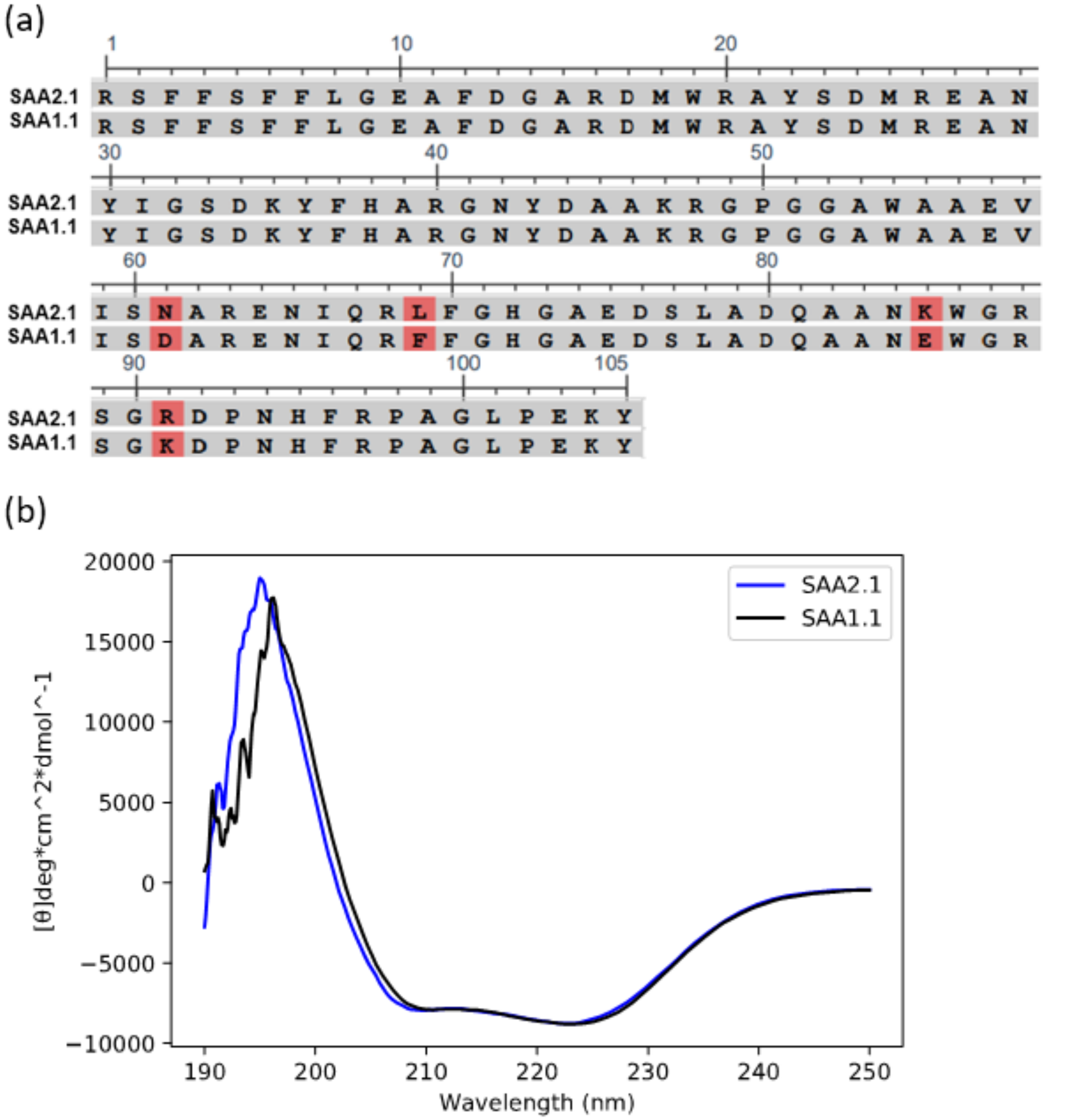


**Figure S1.** (a) Sequence comparison of the two acute phase SAA isoforms SAA2.1 and SAA1.1. Differences are highlighted in red. (b) Circular dichroism spectra of 20µM lipid-free SAA2.1 and SAA1.1 in 20mM Tris (pH 8) at 25 °C.

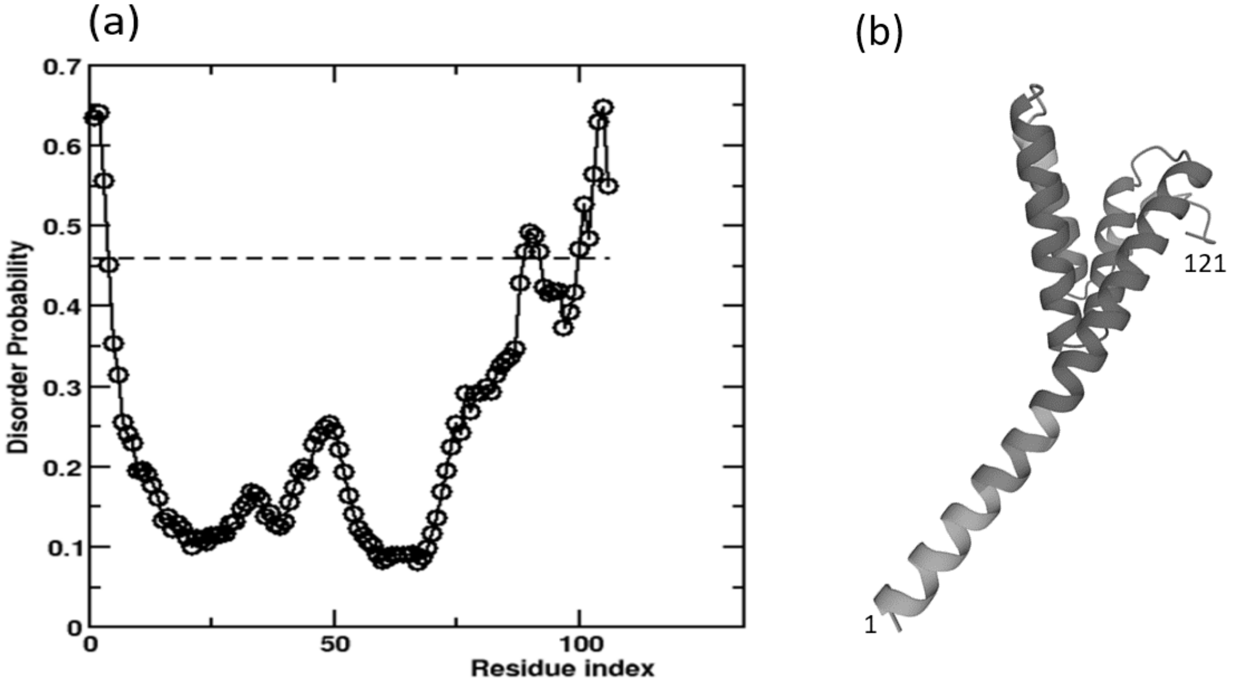


**Figure S2.** (A) Disorder prediction based on machine learning in lipid-free serum amyloid A (106 aa) using MetaPrDOS. Residues beyond the dotted line (threshold) are predicted to be disordered. (B) Secondary and tertiary structure prediction of murine serum amyloid A-1 protein by alphafold (Jumper et al. 2021).

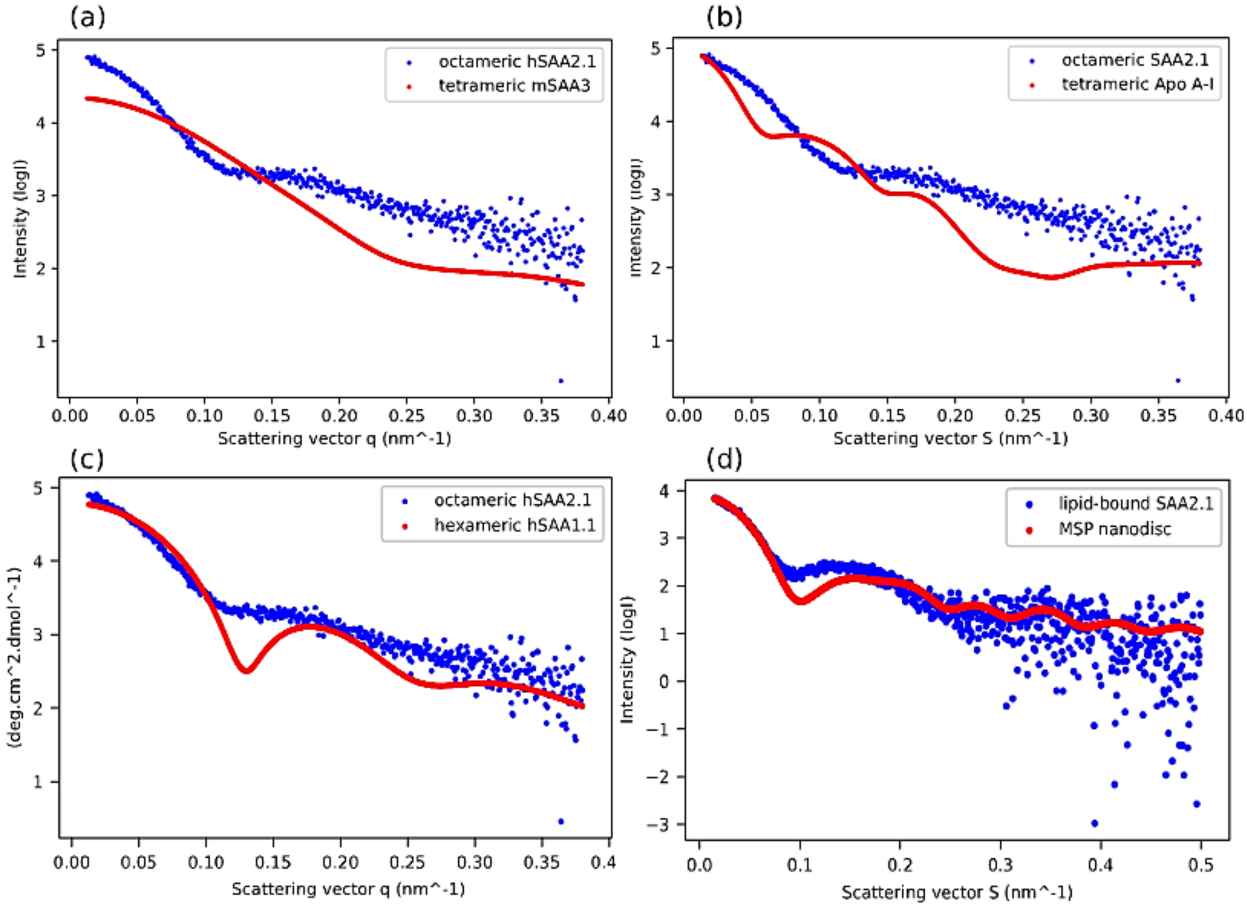


**Figure S3.** (A) Comparison of tetrameric mSAA3 (red, PDB: 4Q5G) and octameric hSAA2.1 in solution (blue) using CRYSOL (ATSAS2.6 package, EMBL, Hamburg, Germany). (B) Comparison of tetrameric Apo A-I (red, PDB: 1AV1) and octameric hSAA2.1 in solution (blue) using CRYSOL (ATSAS2.6 package, EMBL, Hamburg, Germany). (C) CRYSOL output comparing experimental data to hexameric hSAA1.1 (PDB: 4IP9). (D) the CRYSOL output comparing the scattering curves of lipid-bound SAA2.1 and MSP nanodiscs ( $\chi^2$ =0.407, PDB: 2MSC).

(a)

(b)

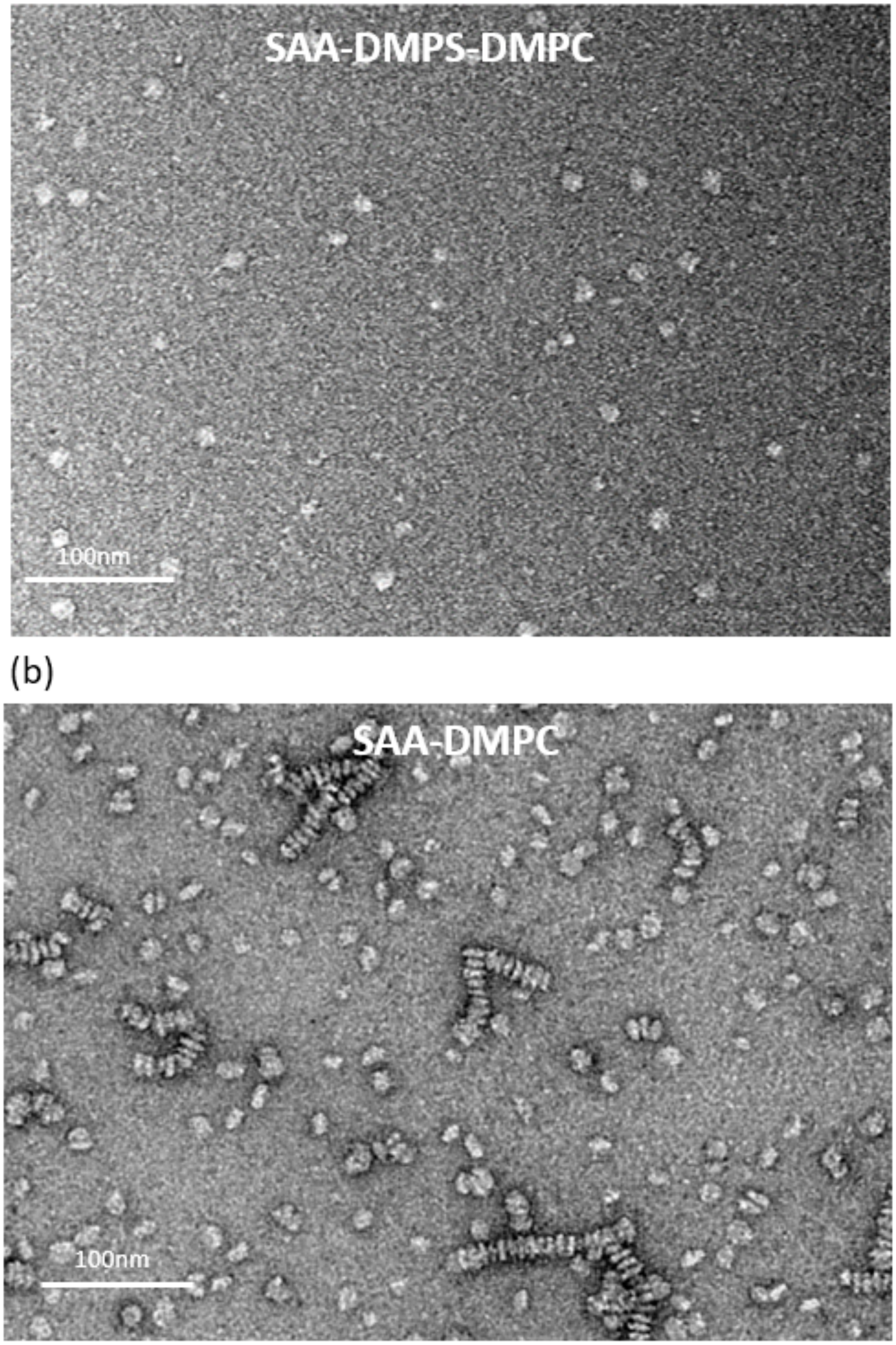


**Figure S4.** (a)Transmission electron microscopy of DMPS-DMPC(80:20)-bound hSAA2.1 and (b) DMPC-bound hSAA2.1 stained with 2% (w/v) uranyl acetate. Nanodiscs were prepared by incubation of 1:1 phospholipid-to-SAA2.1 (w/w) for 2 hrs at 26°C.

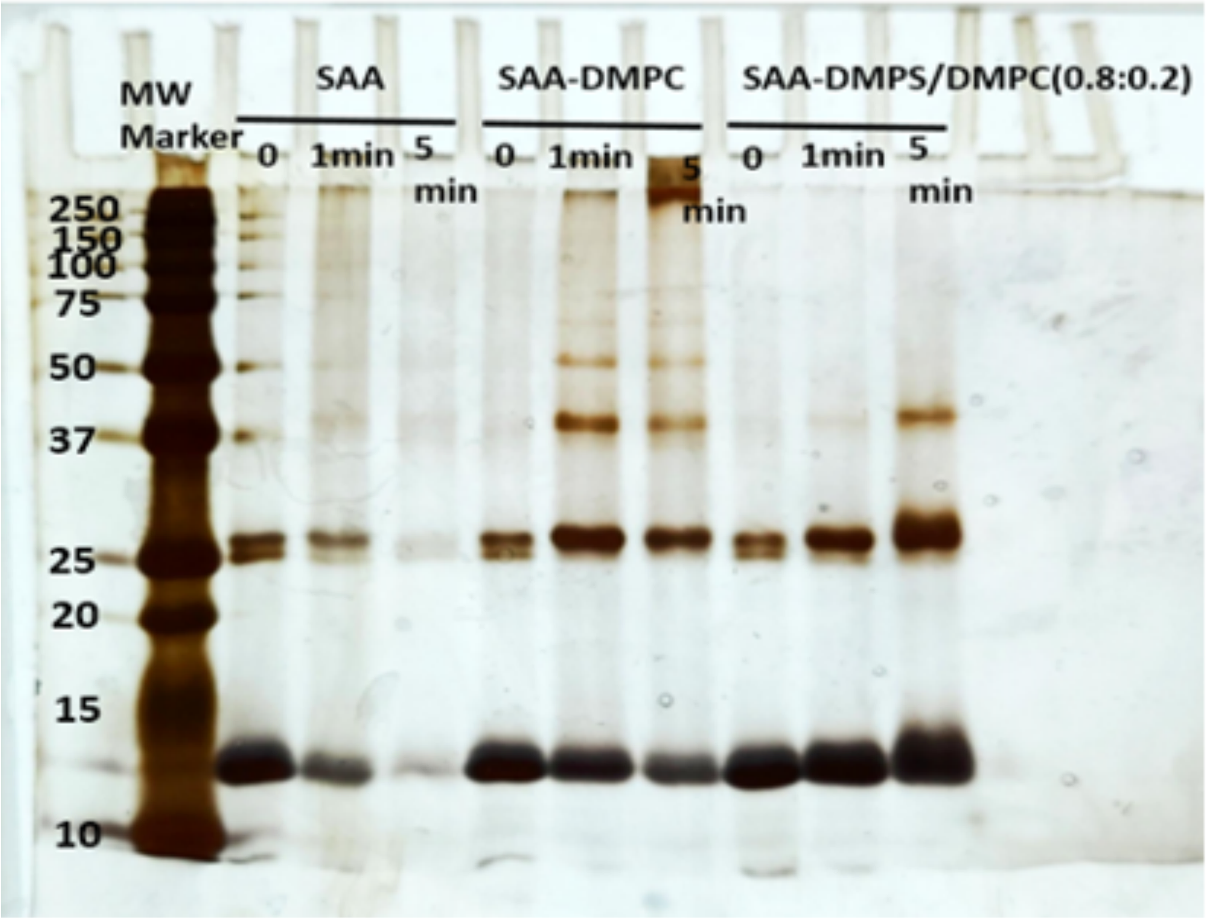


**Figure S5.** Glutaraldehyde crosslinking over time (minutes) followed by SDS-PAGE and silver staining on lipid free, DMPC-bound, and DMPS-DMPC(80:20)-bound SAA.

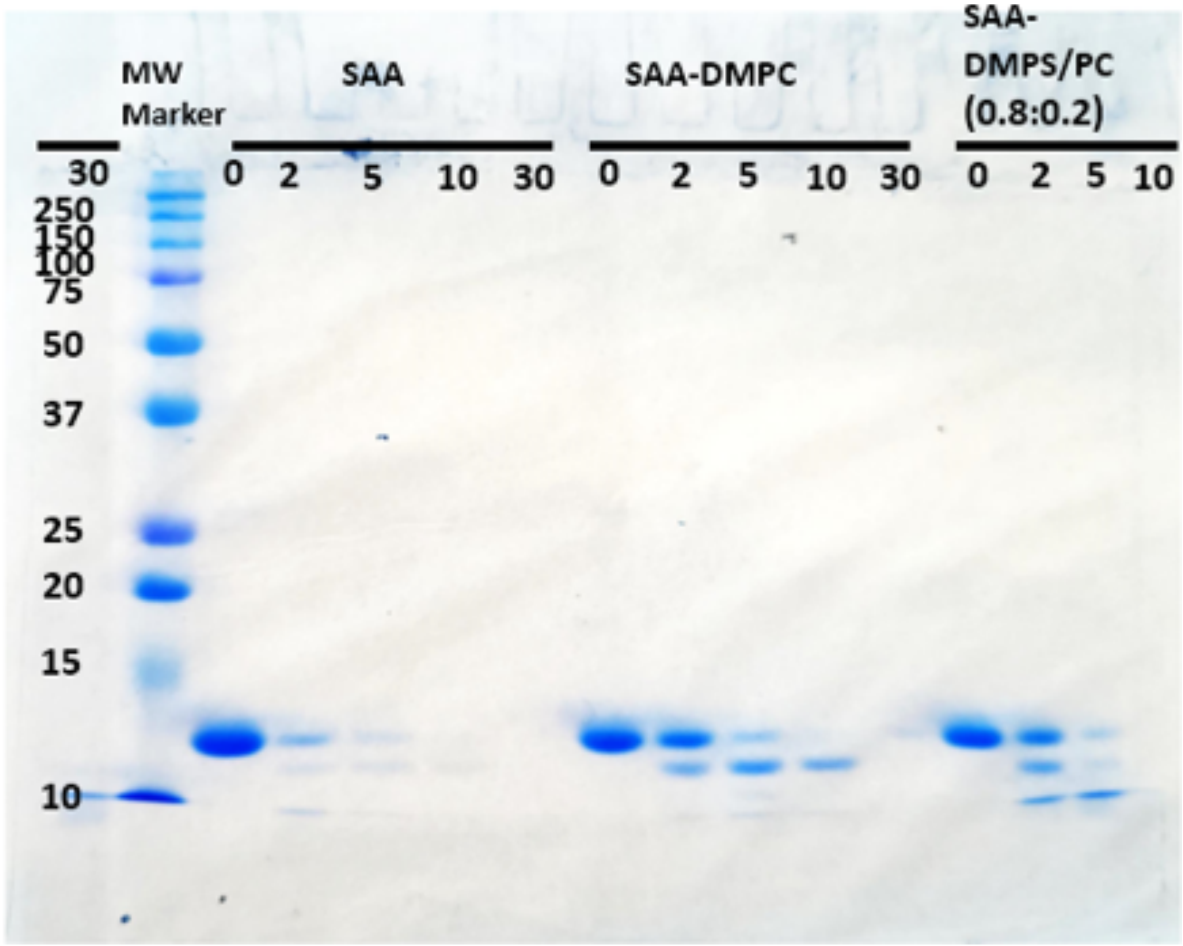


**Figure S6.** Trypsin digestion over time (minutes) followed by SDS-PAGE and Coomassie staining on lipid free, DMPC-bound, and DMPS-DMPC(80:20)-bound SAA.

**Table S1.** Structural parameters of SAA2.1-DMPS/DMPC(80:20) nanodiscs estimated by the *WillItFit* software.

| | |
|---|---|
| **Chi-squared** | **2.51** |
| **Number of Lipids** | **60** |
| **Height of Belt** | **2 nm** |
| **Height of Bilayer** | **3.2 nm** |
| **Height of Hydrophobic Bilayer** | **2.6 nm** |
| **Mol. Volume of Belt** | **41117.2** |
| **Volume of Core** | **36918.6** |
| **Width of Belt** | **1.2 nm** |
| **Volume of Methyl** | **7192.98** |
| **Volume of Headgroups** | **21128.5** |
